# Microtubule binding of the kinesin-4 KIF7 and its regulation by autoinhibition

**DOI:** 10.1101/772327

**Authors:** T. Lynne Blasius, Yang Yue, Kristen Verhey

## Abstract

KIF7 is a member of the kinesin-4 family and plays critical roles in Hedgehog signaling in vertebrate cells. KIF7 is an atypical kinesin as it binds to microtubules but is immotile. We demonstrate that, like conventional kinesins, KIF7 is regulated by autoinhibition as the full-length motor cannot bind to microtubules whereas truncated versions bind statically to microtubules in cells. Previous work suggested that truncated KIF7 motors bind preferentially to the plus ends of microtubules *in vitro*, however, we find that truncated KIF7 does not bind preferentially to or track the plus ends of growing microtubules in mammalian cells or in cell extracts. Although the truncated KIF7 did alter microtubule dynamics in cells, this property is not specific to KIF7 as expression of an active kinesin-1 motor also altered microtubule growth rates. The immotile behavior of KIF7 is not due to the extended neck linker domain as its deletion does not activate KIF7 for motility and its presence in a KIF5C/KIF7 chimeric motor does not prevent processive motility. Together this work indicates that the atypical kinesin KIF7 is regulated by autoinhibition to prevent binding to microtubules and alteration of microtubule dynamics in cells.

## Introduction

Microtubule-based motors of the kinesin superfamily play essential roles in cell division, cell motility, intracellular trafficking, control of microtubule dynamics, and ciliary function [1]. Kinesins are defined by the presence of a ^~^350 aa kinesin motor domain which contains signature sequences for ATP and microtubule binding. Sequence differences within this core motor domain provide family-specific microtubule-based properties. The “conventional” kinesin property of ATP-dependent processive motility along the microtubule surface is a characteristic feature of members of the kinesin-1, kinesin-2 and kinesin-3 families, with sequence changes providing family-specific outputs in terms of speed, processivity, and force generation [2–10]. Unconventional kinesin properties include the sliding of microtubules by kinesin-5 motors and the regulation of microtubule dynamics by members of the kinesin-4, kinesin-8, and kinesin-13 families [11–16].

KIF7 is a member of the kinesin-4 family and plays critical roles in Hedgehog signaling in vertebrates. In mice, knockout of *Kif7* is perinatal lethal with early embryonic defects including preaxial polydactyly, exencephaly, and microphthalmia [17, 18]. In humans, mutations in *KIF7* have been shown to cause severe ciliopathies including Joubert, hydrolethalus, acrocallosal, Meckel–Gruber, and Bardet– Biedl syndromes [19–28]. Recent work has suggested that KIF7 also plays a role during cell proliferation during development and disease [29–31].

In the vertebrate Hedgehog signaling pathway, KIF7 plays both positive and negative roles in Hedgehog signaling by regulating the abundance of Gli2 and Gli3 transcription factors as well as the balance between their activator and repressor forms [17, 18, 32]. In mice, KIF7 activity depends on the presence of the primary cilium and in response to Hedgehog pathway activation, KIF7 localizes to the tip of the primary cilium and facilitates the localization of Hedgehog effectors to the same location [17, 18, 32]. KIF7 has also been suggested to regulate the length of the primary cilium and organization of the cilium tip [33]. Indeed, humans and mice with *Kif7* mutations have longer cilia [23, 33–35]. These ciliary functions of KIF7 are thought to be due to the ability of the motor to bind selectively to the plus ends of microtubules and regulate microtubule dynamics [33, 36]. However, this model is based on *in vitro* characterization of KIF7’s microtubule-based activities and whether KIF7 regulates microtubule dynamics in cells has not been demonstrated.

As a kinesin, KIF7 has several unique properties. Most striking is the fact that KIF7 is not capable of processive motility but rather interacts statically with the microtubule [33, 37]. The mechanistic basis of this immotile behavior is that binding to microtubules does not cause a significant change in the motor’s ATPase activity [36, 37]. Furthermore, KIF7 binds with high affinity to microtubules regardless of its nucleotide state. Still unclear is how the microtubule binding of KIF7 is regulated and how KIF7’s immotile behavior relates to its functions in Hedgehog signaling. Kinesin activity must be tightly regulated in cells to prevent non-productive ATP hydrolysis and misregulation of microtubule-based activities. A general model in the field is that kinesin proteins are regulated by an autoinhibition mechanism where the non-motor regions of the protein act in cis to block the ability of the motor domain to bind ATP and/or microtubules [38]. Here we investigate whether full-length KIF7 motors are regulated by autoinhibition. We also test whether mutation or other sequence changes to the KIF7 motor domain can alter its ability to interact with microtubules and/or undergo processive motion.

## Results

### Full-length KIF7 is autoinhibited

To test whether KIF7 is regulated by autoinhibition, we compared the localization of full-length mouse KIF7 to that of a previously-studied truncated version, KIF7(1-558), which binds to microtubules but is immotile in single-molecule motility assays [33, 37]. Both proteins were tagged with the fluorescent protein monomeric Citrine (mCit). The truncated KIF7(1-558)-mCit motor decorated microtubules in COS-7 cells (Figure 1B, bottom row), consistent with its ability to bind to microtubules in *in vitro* assays [33, 37]. In contrast, the majority of full-length motor protein accumulated in puncta of various sizes scattered throughout the cytoplasm with some protein remaining cytosolic and diffuse (Figure 1B, top row). These puncta may be aggregates or other accumulations of the full-length protein but are not related to the mCit tag as KIF7 tagged with small epitope tags also localized to puncta throughout the cytoplasm (manuscript in preparation). Immunostaining indicates that endogenous KIF7 also localizes to puncta in vertebrate cells [33, 39, 40]. The inability of full-length KIF7 to bind to microtubules suggests that the motor is regulated by autoinhibition.

**Figure 1.**
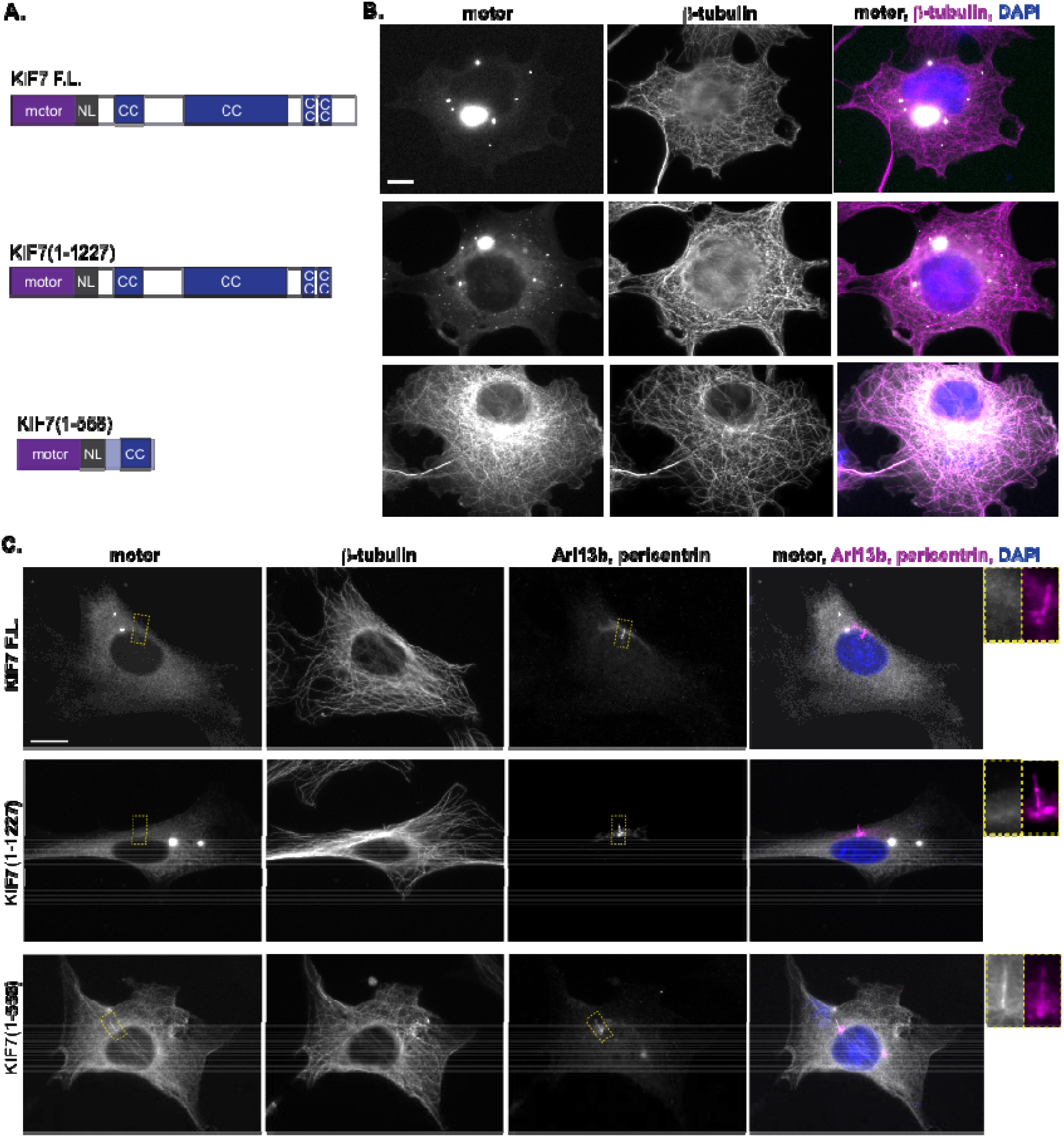
KIF7 is regulated by autoinhibition. (A) Domain organization of full-length (F.L.) KIF7 and the indicated truncation constructs. NL = neck linker. CC = coiled coil. (B) COS-7 cells expressing mCit-tagged FL or truncated versions of KIF7 were fixed and stained with an antibody to β-tubulin to mark microtubules and with DAPI. (C) NIH-3T3 cells expressing mCit-tagged FL or truncated versions of KIF7 were fixed and stained with antibodies to β-tubulin to mark microtubules, Arl13b to mark the primary cilium, pericentrin to mark the basal body, and DAPI to mark the nucleus. Yellow dotted lines indicate boxed regions displayed on far right with magnified views of the primary cilium. Scale bars, 10 μm.

To verify that the full-length and truncated motors are immotile in cells, we expressed the constructs in neuronal cells where the microtubule minus and plus ends are spatially segregated to the cell body and neurite tips, respectively. The ability of active motors to accumulate at neurite tips correlates with their processive motility in single-molecule *in vitro* assays [41–43]. When expressed in neuronal CAD cells, full-length KIF7 localized diffusely throughout the cell and to puncta in the cell body (Figure S1A) whereas truncated KIF7(1-558) bound to microtubules throughout the cell body and proximal region of the neurite (Figure S1B). Neither protein accumulated in the neurite tips, suggesting that neither the full-length nor the truncated motor is capable of utilizing processive motility to reach the distal neurite.

Domain analysis of the KIF7 sequence shows an N-terminal motor domain followed by several regions of potential coiled-coil (CC) and a C-terminal tail domain (Figure 1A). To test whether the C-terminal tail domain of KIF7 is involved in autoinhibition, we created a truncated version KIF7(1-1227) and expressed this construct in COS-7 cells. Similar to the full-length protein, the truncated motor KIF7(1-1227) accumulated in puncta of various sizes scattered throughout the cytoplasm while some protein remained cytosolic and diffuse (Figure 1B, middle row). We conclude that the central region of KIF7 encompassing residues 559-1227, potentially a coiled-coil segment, is involved in autoinhibition of the KIF7 motor domain.

To examine the activity of full-length and truncated versions of KIF7 in a cell line capable of Hedgehog signaling, we expressed these constructs in NIH-3T3 cells. Both the full-length and KIF7(1-1227) constructs displayed a diffuse localization and accumulation in bright puncta (Figure 1C, top and middle rows). In general, the puncta in NIH-3T3 cells appeared smaller than those in COS-7 cells, suggesting that the cellular environment influences the aggregation of these proteins. Under steady-state conditions (cilium maintenance in the absence of Hedgehog stimulation), the full-length motor and the truncated KIF7(1-1227) motor were largely absent from the primary cilium (Figure 1C). When expressed in NIH-3T3 cells, the truncated KIF7(1-558) motor decorated all microtubules including the axoneme in the cilium shaft (Figure 1C, bottom row). In general, the expressed KIF7(1-558) protein maintained more of a cytosolic pool in NIH-3T3 cells as compared to the near-total microtubule binding observed in COS-7 cells, again suggesting that differences in cellular environment can shift motor behavior in these assays. Together these results indicate that full-length KIF7 is autoinhibited whereas the truncated KIF7(1-558) protein has lost this regulation and can interact with all microtubules in cells.

### Truncated KIF7 does not track the plus ends of microtubules in cells

Previous work using purified KIF7(1-560) or KIF7(1-543) in *in vitro* assays suggested that this motor has a higher affinity for GTP-bound tubulin and thus binds preferentially to and tracks with the growing plus ends of microtubules [33, 36]. However, when examined in fixed COS-7, NIH-3T3 or CAD cells, KIF7(1-558) bound to the surface of all microtubules and showed no preference for any subset of microtubules or region such as the plus ends (Figure 1B,C, Figure S1B). We thus used live-cell imaging to examine whether KIF7(1-558) could track the growing ends of microtubules in COS-7 cells.

KIF7(1-558) was tagged with monomeric NeonGreen (mNG) and co-expressed with EB3-mCherry which marks the growing plus ends of microtubules. In live cells, KIF7(1-558)-mNG fluorescence was observed along the shaft of all microtubules (Figure 2A). Although we cannot rule out the possibility that individual KIF7(1-558)-mNG motors bound to the plus ends of microtubules, we did not observe significant or preferential interactions of KIF7(1-558)-mNG with this region (Figure 2A). Despite not binding to the microtubule plus ends, expression of KIF7(1-558)-mNG caused a reduction in the microtubule growth rate to 18.30 ± 0.42 μm/min as compared to control cells expressing mNG (24.18 ± 0.65 μm/min) (Figure 2B,C). To compare these results to that of another motor that is not known to influence microtubule dynamics in cells, we expressed a truncated and active version of the kinesin-1 motor KIF5C(1-559)-mNG. This motor binds preferentially to stable microtubules marked by specific posttranslational modifications ([3], Figure 2A) yet its expression also resulted in a reduction of the average microtubule growth rate to 17.33 ± 0.51 μm/min (Figure 2B,C). Thus, it may be that expression of kinesin motor domains in cells alters microtubule growth rates through indirect means.

**Figure 2.**
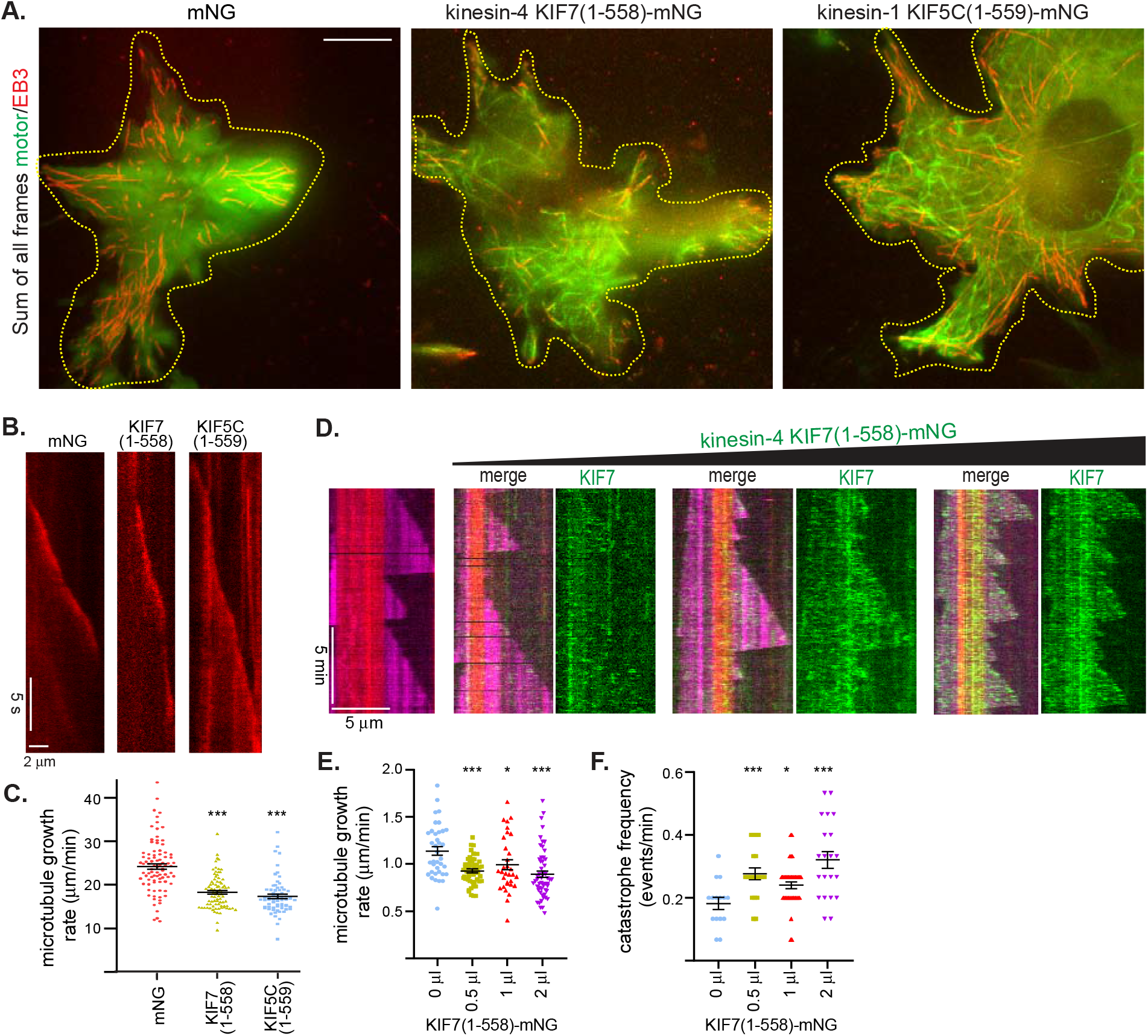
KIF7 does not track the plus ends of microtubules in cells. (A-C) Microtubule dynamics in cells. COS-7 cells expressing EB3-mCherry together with (left panel) mNG control protein, (middle panel) KIF7(1-558)-mNG, or (right panel) KIF5C(1-559)-mNG were imaged live. (A) Representative images show the maximum intensity projection of all frames. Transfected cells are indicated by yellow dotted lines. Scale bar: 10 μm. (B) Representative kymographs of EB3-mCherry in live COS-7 cells expressing mNG (control), KIF7(1-558)-mNG or KIF5C(1-559)-mNG. Time is on the y axis (scale bar, 5 sec); distance is on the x axis (scale bar, 2 μm). (C) Quantification of growth rates of individual microtubules in cells expressing mNG (control), KIF7(1-558)-mNG, or KIF5C(1-559)-mNG. N=66-90 microtubules across 9-10 cells. ***, P<0.001 as compared to control cells (two-tailed *t* test). (D-F) Microtubule dynamics *in vitro*. (D) Representative kymographs of microtubule dynamics in the absence or with increasing amounts of cell lysate containing KIF7(1-558)-mNG. Time is on the y axis (scale bar, 5 min); distance is on the x axis (scale bar, 5 μm). Red= GMPCPP-containing microtubule seeds, magenta= growing microtubules, and green=KIF7(1-558)-mNG. Quantification of microtubule (E) growth rates (N=30-60 events) and (F) catastrophe frequency (N=15-40 microtubules) in the absence or presence of increasing amounts of cell lysate containing KIF7(1-558)-mNG. *, P<0.05 and ***, P<0.001 as compared to control (two-tailed *t* test).

One possible explanation for the differences between the previous work and our results is that the previous work used KIF7 constructs expressed in *E coli* or insect cells whereas we have utilized KIF7 constructs expressed in mammalian cells. We thus tried to replicate the previous *in vitro* work using mammalian-expressed KIF7(1-558) in microtubule dynamics assays. Lysates of cell expressing KIF7(1-558)-mNG were added to flow chambers containing biotinylated microtubule seeds bound to the glass surface together with soluble tubulin for microtubule polymerization. Addition of increasing amounts of KIF7(1-558)-mNG resulted in a decrease in the microtubule growth rate and an increase in the microtubule catastrophe rate (Figure 2D), consistent with previous work using KIF7(1-558) expressed in *E coli* or insect cells [33, 36, 37]. However, we did not observe the mammalian-expressed protein to bind preferentially to either the GMPCPP-containing microtubule seed or the GTP-containing microtubule plus ends (Figure 2D).

### Mutation of residues that coordinate nucleotide alters the microtubule binding of KIF7

Although KIF7 shows little to no microtubule-stimulated ATPase activity [37], recent work using recombinant proteins suggested that the nucleotide state of KIF7 can determine its preferential binding to the plus end of microtubules [36]. Specifically, in the presence of ATP, ADP or ATPγS, KIF7(1-543) was found to bind preferentially to the growing plus ends of microtubules whereas in the presence of the non-hydrolysable analog AMPPNP, KIF7(1-543) was found to bind along the lattice of the microtubule [33, 36]. We thus tested whether mutations in the nucleotide-binding pocket could alter the ability of KIF7 to bind preferentially to the plus ends of microtubules in cells.

The P-loop motif [GxxxxGK(S/T)] is conserved across the kinesin superfamily and required for coordination of Mg^2+^ and nucleotide (Figure S2B). In kinesin-1, mutation of the last residue of the P loop, T93, to Asparagine results in a motor that is locked in the ATP-like state and binds strongly to microtubules (a “rigor” mutant) [44]. We thus generated a KIF7(1-558) construct containing the analogous mutation, T101N (Figure 3A, Figure S2). While generating this construct, we also identified a construct with a point mutation, introduced during PCR, that changed the F86 residue to leucine (F86L, Figure 3A, Figure S2). As this residue is a Tyrosine in all other kinesin-4 motors as well as in kinesin-1 motors (Figure S2), we analyzed the microtubule-binding capacity of the KIF7(1-558, F86L) mutant as well.

**Figure 3.**
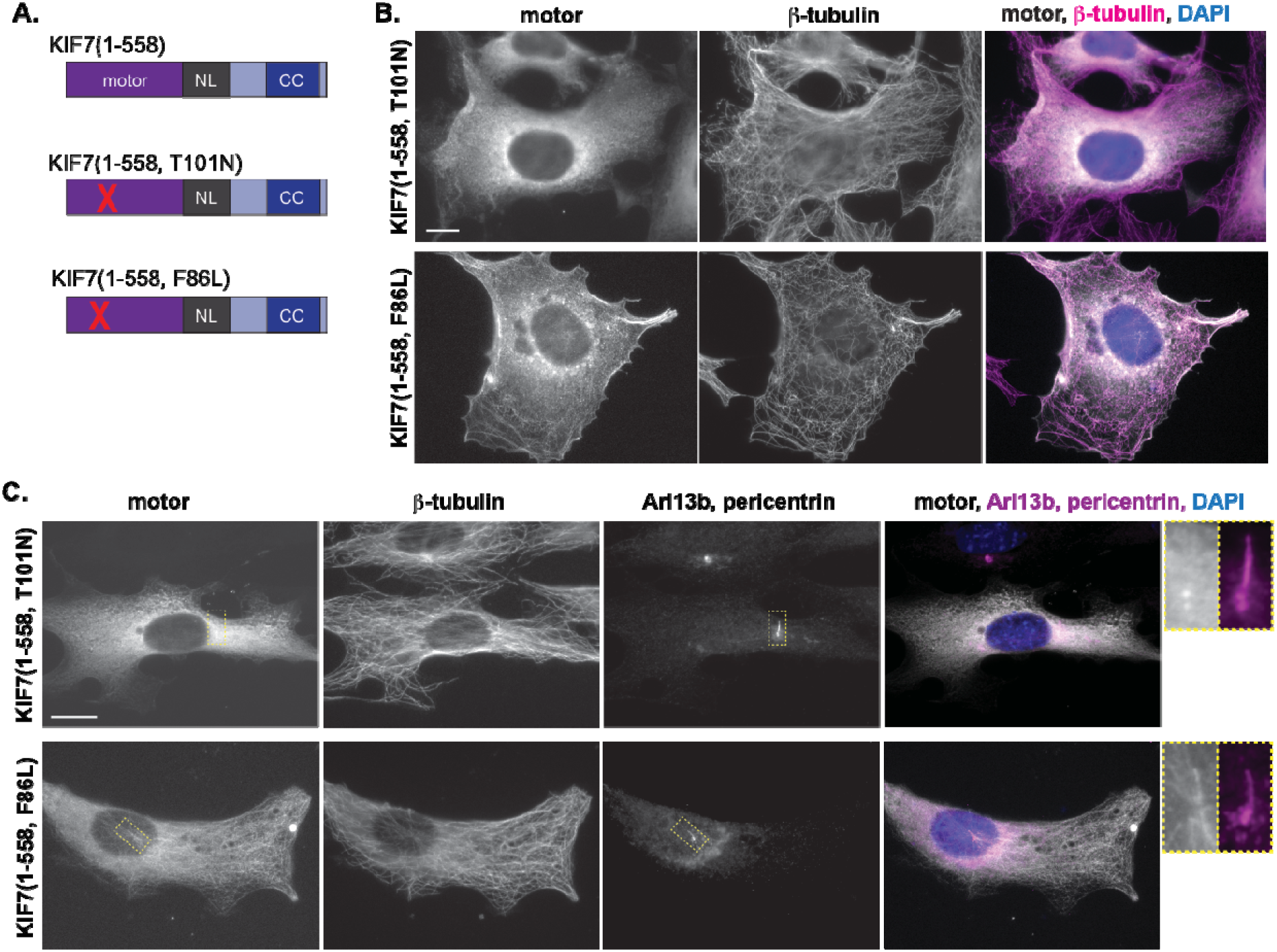
Mutations in or near the nucleotide-binding pocket of KIF7 can alter microtubule binding. (A) Domain organization of the indicated truncated KIF7 motor constructs. Red X indicates location of point mutations. (B) COS-7 cells expressing the indicated mCit-tagged versions of KIF7 were fixed and stained with an antibody to β-tubulin to mark microtubules and with DAPI. (C) NIH-3T3 cells expressing the indicated mCit-tagged versions of KIF7 were fixed and stained with antibodies to β-tubulin to mark microtubules, Arl13b to mark the primary cilium, pericentrin to mark the basal body, and with DAPI. Scale bars, 10 μm. Yellow dotted lines indicate boxed regions displayed on far right with magnified views of the primary cilium.

When expressed in COS-7 cells, KIF7(1-558, T101N) did not cause the motor to bind to microtubules in a “rigor” manner as expected. Rather, the expressed motor localized in a diffuse and cytosolic appearance (Figure 3B), suggesting that this mutation abolished the ability of KIF7(1-558) to bind to microtubules. Similar results were obtained upon expression of the KIF7(1-558, T101N) construct in NIH-3T3 cells (Figure 3C). In contrast, mutation of F86L had no effect on the ability of KIF7(1-558) to bind to microtubules in COS-7 (Figure 3B) or NIH-3T3 cells (Figure 3C). Using a fluorescence-based *in vitro* microtubule binding assay, we quantified the relative microtubule affinities of the mutant motors and determined that mutation of F86L did not alter the affinity of KIF7 for microtubules in the presence of ATP whereas mutation of T101N caused a >5-fold decrease in the affinity of KIF7 for microtubules (Figure S2C).

### Replacement of surface loops of KIF7 can alter microtubule binding

KIF7 shares 37% sequence identity in the motor domain with the kinesin-1 motor KIF5B and sequences specific to KIF7 are likely responsible for decoupling the chemical and mechanical states of this motor [45]. Most sequence differences with KIF7 are found in the surface loops that connect secondary structure elements (Figure S2). To test whether the sequence alterations in these loops influence the ability of KIF7 to bind microtubules and/or couple microtubule binding to motility, we created swap constructs where the loop sequences of KIF7 were replaced by the analogous sequences from the kinesin-1 motor KIF5C. We focused on sequence changes to loop5 and loop10 as these sequences in KIF7 are different not only from kinesin-1, but also from motile kinesins within the kinesin-4 family (Figure S2B).

When expressed in COS-7 cells, a KIF7(1-558) construct containing the loop 5 (L5) sequences of KIF5C [Figure 4A, KIF7(1-558, swap L5)] bound to all microtubules (Figure 4B) suggesting that replacement of L5 did not diminish, and perhaps enhanced, the microtubule binding capacity of KIF7. Similar results were obtained upon expression of the swap L5 construct in NIH-3T3 cells as the KIF7(1-558, swap L5) motor bound to all microtubules including the axoneme of the primary cilium (Figure 4C), again suggesting that replacement of L5 did not alter the microtubule binding of KIF7. In contrast, replacement of loop 10 (L10) of KIF7 with the analogous sequences of KIF5C [Figure 4A, KIF7(1-558, swap L10)] resulted in a motor that was cytosolic and diffuse in COS-7 (Figure 4B) and NIH-3T3 (Figure 4C) cells. This motor failed to bind not only to cytoplasmic microtubules but also to axonemal microtubules of the primary cilium (Figure 4C). These results suggest that replacement of L10 abolished the ability of KIF7(1-558) to bind to microtubules in cells.

**Figure 4.**
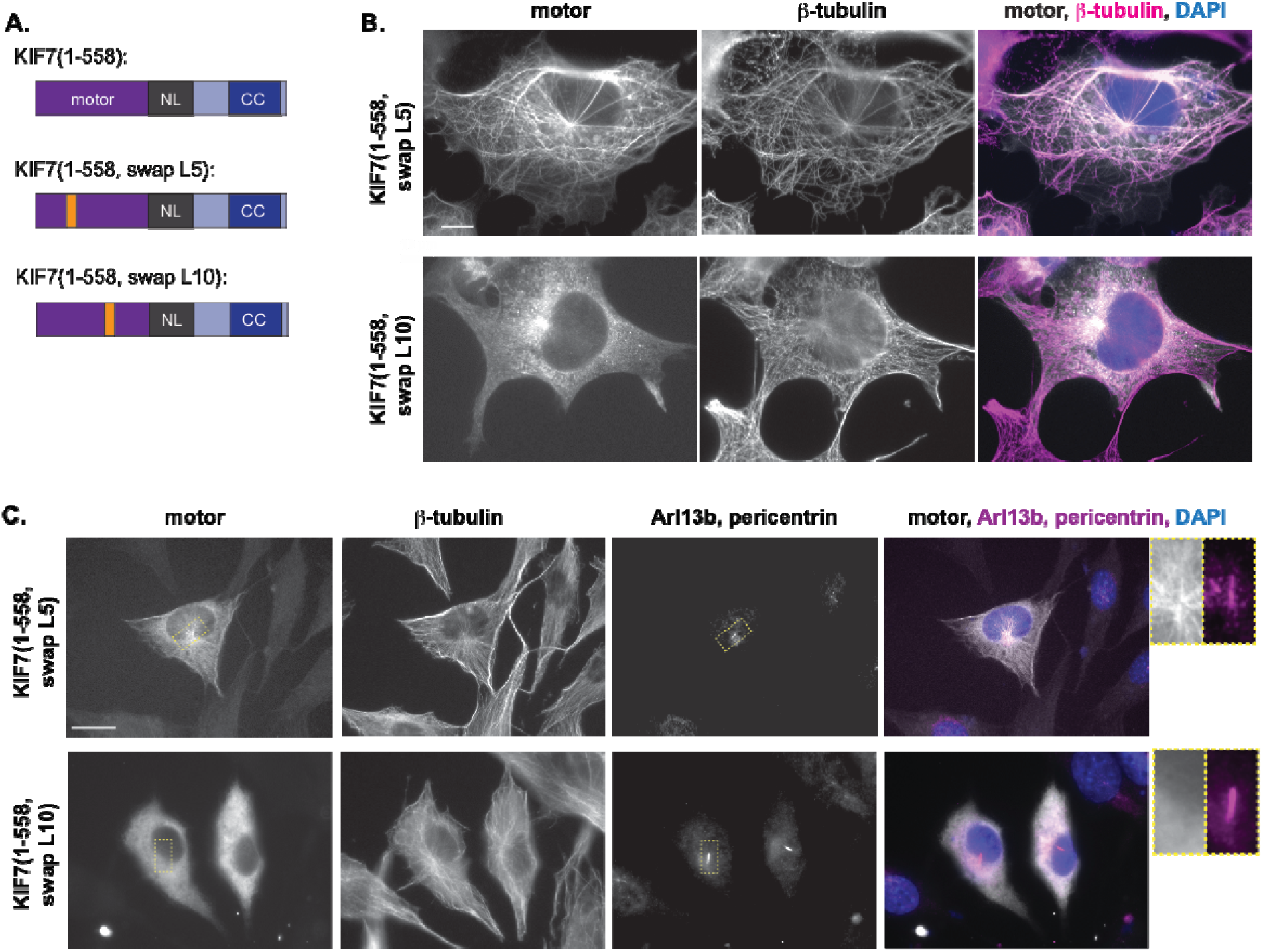
Replacing loop 10 reduces microtubule binding whereas replacing loop 5 enhances microtubule binding. (A) Domain organization of the indicated truncated KIF7 motor constructs. Orange boxes indicate the replacement of the loop 5 (L5) or loop 10 (L10) regions of KIF7 by those of the kinesin-1 KIF5C. (B) COS-7 cells expressing the indicated mCit-tagged versions of KIF7 were fixed and the indicated mCit-tagged versions of KIF7 were fixed and stained with antibodies to β-tubulin to mark microtubules, Arl13b to mark the primary cilium, pericentrin to mark the basal body, and with DAPI. Scale bars, 10 μm. Yellow dotted lines indicate boxed regions displayed to the far right with magnified views of the primary cilium.

### The extended neck linker of KIF7 does not prevent microtubule-based motility

An unusual feature of the KIF7 domain structure is that the neck linker (NL, a structural element that coordinates the two motor domains in motile kinesins) is separated from the first coiled-coil (CC, required for dimerization in processive kinesins) by an extended region of ^~^120 amino acids (Figure 1A, Figure S3A,E). This extended NL could prevent coordination of the two motor domains in a dimeric KIF7 motor and could therefore be responsible for the lack of motility. To test this, we generated two deletion constructs, Δ357-509 and Δ362-488, designed to remove the extended NL and generate a NL-to-CC transition similar to that of processive kinesins (Figure 5A, Figure S3E,F). When expressed in COS-7 cells, KIF7(1-558,Δ357-509) and KIF7(1-558,Δ362-488) constructs did not bind to microtubules but rather displayed a diffuse cytosolic localization (Figure 5B). Similar results were obtained upon expression of the KIF7(1-558, Δ357-509) and KIF7(1-558, Δ362-488) constructs in NIH-3T3 cells (Figure 5C). Furthermore, these constructs were not capable of processive motility as they did not accumulate at the plus ends of microtubules in neurites of CAD cells (Figure S1C,D). These results suggest that rather than activate KIF7(1-558) for motility, deletion of the extended NL abolished the ability of the motor to bind to the microtubule lattice.

**Figure 5.**
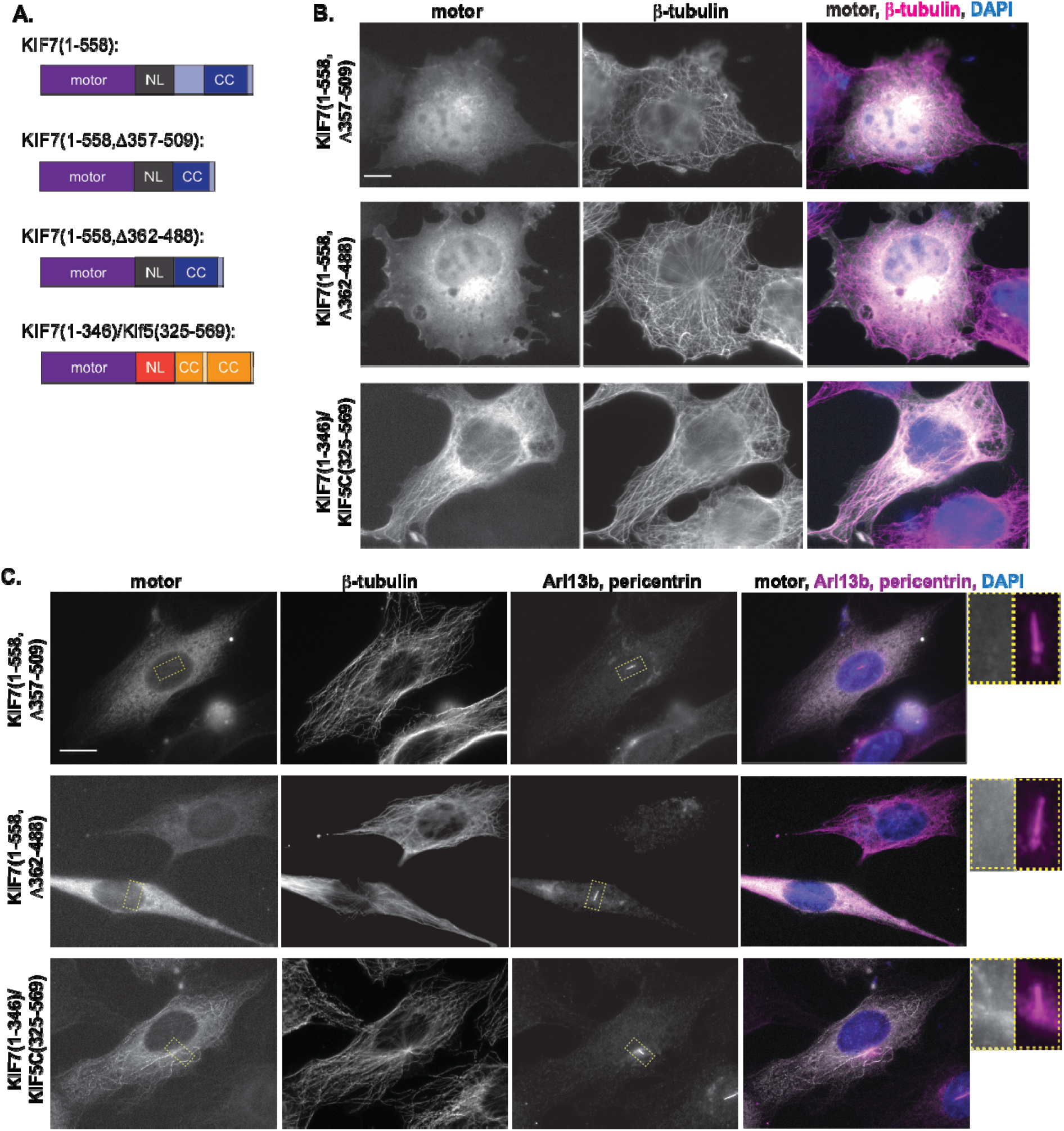
Removal of the extended NL does not activate KIF7 for motility. (A) Domain organization of the indicated truncation or chimeric motor constructs. (B) COS-7 cells expressing mCit-tagged truncated or chimeric versions of KIF7 were fixed and stained with an antibody to β-tubulin to mark microtubules and with DAPI. (C) NIH-3T3 cells expressing mCit-tagged FL or truncated versions of KIF7 were fixed and stained with antibodies to β-tubulin to mark microtubules, Arl13b to mark the primary cilium, pericentrin to mark the basal body, and with DAPI. Scale bars, 10 μm. Yellow dotted lines indicate boxed regions displayed on far right with magnified views of the primary cilium.

As an alternative approach to removing the extended NL of KIF7, we replaced the NL and CC segments of KIF7 with those of the processive kinesin-1 motor KIF5C [Figure 5A, construct KIF7(1-346)/KIF5C(325-569), Figure S3F]. Fusion to the first CC of kinesin-1 has been used previously to examine the motility properties of kinesin-2 and kinesin-3 motors [46, 47]. The chimeric motor KIF7(1-346)/KIF5C(325-569) behaved similar to the KIF7(1-558) construct and bound to microtubules in both COS-7 (Figure 5B) and NIH-3T3 cells (Figure 5C). Although this construct was able to bind to microtubules, it was not capable of processive motility as it failed to accumulate at the plus ends of microtubules in the neurites of CAD cells (Figure S1E). Thus, engineering a NL-to-NC transition that is compatible with motility in other kinesin motors did not generate a KIF7 motor capable of chemomechanical coupling and motility. This result suggests that the defect in KIF7 motility is within the core motor domain itself rather than the coupling of the two motor domains in a dimeric molecule.

### KIF7 becomes an active motor when fused to the motor domain of kinesin-1

Although the immotile behavior of KIF7 appears to be due to sequences specific to the motor domain, it is possible that sequences outside the motor domain also contribute to the inability of this motor to undergo processive motility. To test this, we attempted to generate an active KIF7 motor by replacing the motor and neck linker of KIF7 with the minimum sequences of the kinesin-1 motor KIF5C required for processive motility (motor+NL+CC, Figure 6A). When expressed in COS-7 cells, the motor swap construct KIF5C(1-379)/KIF7(369-1348) appeared to be an active motor as the KIF7-containing puncta accumulated at the plus ends of the microtubules localized at the cell periphery (Figure 6B, bottom row). This is in contrast to the wild-type KIF7 protein where the puncta accumulate in the middle of the cell (Figure 6B, top row). Similar results were obtained upon expression of wild-type and swap motor constructs in NIH-3T3 cells (Figure 6C). Furthermore, the active KIF5C(1-379)/KIF7(369-1348) protein showed an increased accumulation at the plus ends of microtubules at the tip of the primary cilium (Figure 5C). These results suggest that the non-motor regions of KIF7 do not prevent motility of the kinesin motor domain. These results also suggest that the non-motor regions of KIF7 are sufficient to target a non-ciliary kinesin to the primary cilium.

**Figure 6.**
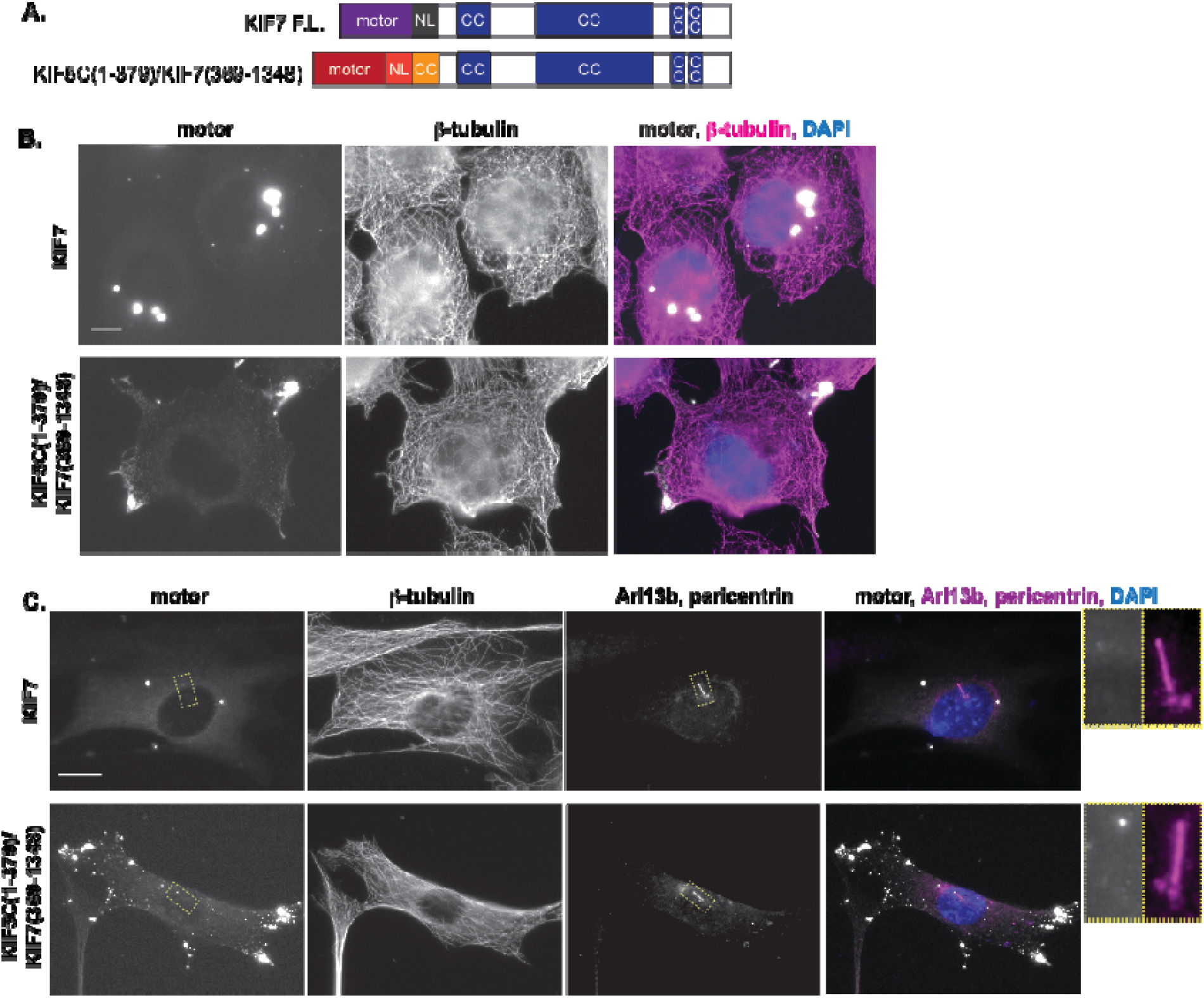
Fusion to the KIF5C motor domain generates an active KIF7 motor. (A) Domain organization of the indicated KIF7 constructs. Red and orange boxes indicate the motor domain, NL, and CC sequences of the kinesin-1 motor KIF5C. (B) COS-7 cells expressing full-length or chimeric mCit-tagged versions of KIF7 were fixed and stained with an antibody to β-tubulin to mark microtubules and with DAPI. (C) NIH-3T3 cells expressing full-length or chimeric mCit-tagged versions of KIF7 were fixed and stained with antibodies to β-tubulin to mark microtubules, Arl13b to mark the primary cilium, pericentrin to mark the basal body, and with DAPI. Scale bar, 10 μm. Yellow dotted lines indicate boxed regions displayed on far right with magnified views of the primary cilium.

## Discussion

### KIF7 is regulated by autoinhibition

A general model for kinesin regulation involves autoinhibition of the motor domain by non-motor regions of the protein. We demonstrate that this model applies to KIF7 as the full-length motor is unable to bind to microtubules whereas deletion of non-motor segments (coiled-coil and tail domains) results in a motor that is competent for microtubule binding in both *in vitro* assays and in cells. Thus, the general model for kinesin regulation extends to family members that are immotile as their microtubule-binding activity is regulated by an autoinhibition mechanism.

Within the kinesin-4 family, the KIF21A motor has previously been demonstrated to be regulated by autoinhibition. For KIF21A, the mechanism of autoinhibition involves direct interactions of the coiled-coil domain with the motor domain. Mutations in this regulatory coiled-coil relieve autoinhibition and lead to the disease congenital fibrosis of the extraocular muscles type 1 (CFEOM1) [48–50]. An analogous regulatory coiled-coil segment plays a role in autoinhibition of KIF21B [51]. As noted by Bianchi et al. [49], the regulatory coiled-coil segment of KIF21A shares 37.5% sequence similarity to amino acids 998-1077 of KIF7, suggesting that the mechanism of autoinhibition for KIF7 may involve interactions between this coiled-coil region and the motor domain. Our work is consistent with this possibility as constructs that retain the coiled-coil region of KIF7 (i.e. 1-1229) are autoinhibited whereas constructs that lack this region (i.e. 1-558) are active for microtubule binding. Interestingly, several disease-associated mutations (N1060S and R1068W [19, 23]) lie in this predicted regulatory domain, suggesting that these residues may cause disease by relieving autoinhibition of KIF7.

### KIF7 does not bind preferentially to or track microtubule plus ends in cells

Previous work using purified proteins and *in vitro* microtubule binding assays demonstrated that KIF7 binds preferentially to GTP-containing microtubule structures such as those in microtubule seeds and plus ends. However, we find that truncated versions of KIF7 do not bind preferentially to or track the plus ends of microtubules in cells, including COS-7 fibroblast-like, NIH-3T3 fibroblast, or CAD neuronal cells. Rather, we find that truncated KIF7(1-558) bind along the surface of all microtubules in cells. Similar to our observations, the previous work of He et al. did observe that KIF7(1-560)-GFP bound along all microtubules, not to the plus ends, and resulted in microtubule bundling in cells [33]. These results highlight the importance of verifying conclusions drawn from *in vitro* assays with experiments carried out in a physiological context.

Why do truncated KIF7 constructs bind preferentially to microtubule plus ends in *in vitro* assays but not in cells? We believe that the most likely explanation is inherent differences between the assays. One possibility is that KIF7’s microtubule binding is highly sensitive to ionic conditions. Support for this possibility can be seen in the *in vitro* work of Jiang et al [36] where monomeric KIF7(1-386)-GFP localized to the plus ends of growing microtubules at 1× BRB80 but along the shaft of growing microtubules at 0.5× BRB80. A second possibility relates to differences between microtubules in cells and those assembled for *in vitro* assays. In cells, microtubules consist of strictly 13-protofilaments and this structural arrangement may influence KIF7 binding. In addition, in cells, numerous proteins localize to and regulate microtubule plus ends and may negatively impact KIF7 at this location. Indeed, the ability of high levels of KIF7(1-543) to destabilize microtubules appeared to be negated by the presence of EB1 in the *in vitro* assay [36]. A third possibility is that the nucleotide state of KIF7 differs between the *in vitro* and cellular assays. The previous studies utilized dimeric KIF7-GFP constructs expressed in *E coli* or insect cells and were found to contain ADP in the nucleotide pocket despite the presence of ATP in the purification buffers [33, 36]. The motors in our study are expressed in mammalian cells and may contain ATP in the nucleotide pocket.

We attempted to drive the localization of KIF7(1-558) to the plus ends of microtubules in cells by introducing a mutation in the nucleotide-binding P-loop. For processive kinesin motors, mutation of the last residue of the P-loop (e.g. Threonine to Asparagine) mimics the apo state and thus results in motors with increased, even “rigor” type, microtubule binding [44]. For KIF7, we previously showed that KIF7(1-558) bound strongly to microtubules under buffer conditions meant to mimic the no nucleotide state [37]. We were thus surprised to find that mutation of the last residue of the P-loop, T101N, abolished the ability of KIF7(1-558) to bind to microtubules in an *in vitro* assay and caused the motor to localize in a diffuse and cytosolic appearance in both COS-7 and NIH-3T3 cells (Figure 5B). Although we cannot rule out the possibility that the T101N mutation altered the folding of the KIF7 motor domain, it is interesting to note that mutation of the analogous residue caused a decrease in microtubule affinity for the *Drosophila* motor Nod (*nod*^*[DTW]*^ mutation), an immotile kinesin that retains microtubule-stimulated ATPase activity perhaps to regulate transient attachments of chromosomes to microtubules rather than to produce transport [52].

### Kinesin-4 motors as regulators of microtubule dynamics

All vertebrate members of the kinesin-4 family (KIF4, KIF21A, KIF21B, KIF7, and KIF27) have been demonstrated to impact microtubule dynamics using purified proteins in *in vitro* assays. Xklp1 (*Xenopus* Kif4A homolog) and Kif21A have been found to decrease microtubule growth rates and suppress catastrophes, generally resulting in microtubule pausing [50, 53]. In contrast, KIF7 and KIF27 decrease the microtubule growth rate and growth length, thus destabilizing microtubules [33, 37]. The influence of KIF21B on microtubule dynamics is unclear as one study suggested that KIF21B increases microtubule growth rates and another suggested that it decreases growth rates [51, 54].

A critical issue is how these *in vitro* results relate to the cellular functions of the kinesins. For KIF4, depletion of the motor results in persistent microtubule growth during mitosis [55–58], consistent with the *in vitro* studies. For KIF21A, its loss of function resulted in an increase in microtubule growth rates whereas overexpression of KIF21A results in decreased microtubule growth [48, 50], also consistent with the *in vitro* results. The physiological relevance of KIF21B’s ability to influence microtubule dynamics are unclear as two studies suggested that KIF21B decreases growth rates and a third study demonstrated an increase in growth rates in cells [51, 54, 59]. For KIF7, loss of KIF7 expression or point mutations in *Kif7* lead to increased cilium length [23, 33–35], consistent with KIF7 functioning to destabilize microtubules. Our results are also consistent with a microtubule destabilizing function for KIF7 in cells as expression of KIF7(1-558) caused a decrease in the microtubule growth rate as measured by EB3-mCherry. However, the ability to influence microtubule dynamics was not unique to KIF7 as overexpression of truncated version of the kinesin-1 motor KIF5C also caused a decrease in the microtubule growth rate. These results suggest that overexpression of active kinesin motors, above physiological levels, can influence microtubule dynamics in cells. This influence is likely to be indirect as neither KIF5C, KIF7 nor KIF21B regulated microtubule dynamics via localization to the plus ends (this study, [51]). One possibility is that the overexpressed motor domains may bind to soluble tubulin and remove subunits from the polymerization-competent pool. Indeed, the KIF7(1-543) motor forms a stable complex with soluble tubulin heterodimers [36].

### Contribution of the extended neck linker and surface loops to KIF7’s immotile behavior

A unique feature of KIF7 is the extended sequence that separates the end of the neck linker and the beginning of the first predicted coiled coil that could drive dimerization. The neck linker is a structural feature required for processive motility of dimeric kinesin motors as it coordinates the catalytic cycles of two motor domains, determines the directionality of stepping, and drives force generation [60] and an extended neck linker was thought to be incompatible with the conventional kinesin stepping mechanism [46, 47, 61]. However, the *Arabidopsis thaliana* motor protein phragmoplast-associated kinesin-related protein 2 (PAKRP2) contains a 32 aa neck linker, yet individual motors are processive with an average velocity of ^~^ 65 nm/s and run length of ^~^3 um [62]. Longer extensions of the neck linker can be found in the kinesin-6 motor subunits of centralspindlin (HsMKLP-1, CeZen-4, DmPav, ^~^70 aa extension) and the kinesin-10 motor KID (HsKIF22, ^~^85 aa extension). For these motors, the initial segment of the neck linker can adopt a backwards-docked conformation that effectively shortens the neck linker [63, 64]. The backwards-docked conformation has been shown to be required for CeZen-4 to drive motility in a multi-motor context [63, 65] and may likewise be required for the ability of KID to function in a similar manner [66, 67].

We tested the possibility that the immotile behavior of KIF7 is caused by its extended neck linker. However, we found that deleting the extended region to align the neck linker and first coiled coil of KIF7 does not result in processive motility, nor does replacing the neck linker and extended region with the neck linker and coiled-coil regions of kinesin-1. While it is possible that our protein engineering did not align the neck linker segments in a manner compatible with interhead tension, these results support the idea that sequence changes within the core motor domain of KIF7 are responsible for this motor’s lack of motility. Consistent with this possibility, recent structural analysis of the monomeric KIF7 motor domain bound to microtubules indicates that several key structural features that alter their conformation in response to nucleotide state in processive kinesins [e.g. L11, α4, and L9 (switch 1)] failed to adopt distinct structural states in ADP and AMPPNP for KIF7 [36].

Our results do not rule out the possibility that a partner protein could bind to KIF7’s extended neck linker to alter the effective length and/or structure in a manner that changes KIF7’s microtubule-based functions. This possibility was suggested by Davies et al [68] based on a) a comparison of kinesin motors with extended neck linkers and b) the finding that the extended neck linker of the kinesin-6 CeZen-4 serves as the binding site for the partner protein CeCyk-4 which alters *Ce*Zen-4 structure to slow motor velocity and enable microtubule bundling [68, 69]. Further support for this idea comes from work on *Ce*Zen-4’s homolog *Dm*Pav where binding of the *Dm*Tum partner enables multi-motor microtubule gliding activity [70]. Notably, the N-terminal half of KIF7 has been found to interact with both Gli2 and Gli3 [18] and the extended neck region has been found to interact with Dlg5 and the C-terminal tail of Smoothened [71]. The effects of these interactions on KIF7’s microtubule-based functions have not been investigated.

We also tested the possibility that the unique surface loops of KIF7 contribute to the immotile behavior of this motor. We found that replacement of L5 with that of kinesin-1 had no effect on microtubule binding or immotile behavior of KIF7. We also found that replacement of L10 with that of kinesin-1 abolished KIF7’s ability to interact with microtubules in cells. This result was surprising as Jiang *et al* [36] recently demonstrated that swapping L10 in HsKIF7 with the corresponding sequence in kinesin-1 led to no change in microtubule affinity based on the behavior of purified proteins in a microtubule co-sedimentation assay. These different results are likely due to differences in the exact swapped sequences as the previous change of *Hs*KIF7’s RGRAPSRLPRPAPGQL to *Hs*KIF5C’s ENVETEKK added two additional Lysine residues as compared to our change of *Mm*1KIF7’s RGRTPSRLPRPAAG to *Hs*KIF5C’s ENVETE. It is also possible that differences between the previous use of *in vitro* assays and our analysis in cells could explain the different behavior of L10 swap motors, including differences in ionic conditions, presence of microtubule associated proteins, and nucleotide state of the expressed motors. Further work is needed to test these possibilities.

## Material and Methods

### Plasmids and antibodies

Plasmids for expression of full-length and (1-558) versions of mouse KIF7 have been described [37]. Additional truncated, mutant or chimeric versions were generated by PCR or gene synthesis and subcloned using appropriate restriction enzymes. All plasmids were verified by DNA sequencing. Antibodies purchased for immunoflouorescence staining: Arl13b (1:1000, Protein Tech Group, 17711-1-AP), Pericentrin (1:500, AbCam, ab4448), β-tubulin (1:5.000 clone E7; Developmental Studies Hybridoma Bank).

### Cell culture, transfection, and immunofluorescence

COS-7 cells (monkey kidney fibroblast, ATCC, RRID:CVCL_0224) and NIH-3T3 cells (ATCC; RRID:CVCL_0594) were cultured in DMEM (Gibco) with 10% (vol/vol) Fetal Clone III (HyClone) and GlutaMAX (Gibco) at 37 °C with 5% CO2. Mouse central nervous system catecholaminergic CAD cells (RRID:CVCL_0199) were grown in a 1:1 mixture of F12:DMEM (Gibco) plus 10% FBS (HyClone) and GlutaMAX (Gibco) at 37 °C with 5% CO2. For NIH-3T3 cells, cells were switched to low serum media at the time of transfection to arrest cells in G1 and fixed and stained 48 h later. For CAD cells, 12 h after transfection, cells were switched to differentiation media (F12:DMEM without serum) to induce neurite outgrowth and 48 h after transfection were fixed and processed for indirect immunofluorescence. Transfections were carried out using Trans-IT LT1 (Mirus) in Opti-MEM (Gibco) according to the manufacturer’s instructions.

COS-7 cells were fixed with 3.7% formaldehyde in PBS, treated with 50 mM NH_4_Cl in PBS to quench unreacted formaldehyde, permeabilized with 0.2% Triton X-100 in PBS, and then blocked in blocking solution (0.2% fish skin gelatin in PBS). NIH-3T3 were fixed with ice cold methanol for 10 minutes, rinsed with PBS, and blocked in blocking solution (0.2% fish skin gelatin in PBS). Primary and secondary antibodies were applied in blocking solution at room temperature for 1 h each. Nuclei were stained with 10.9 μM 4⍰6-diamidino-2-phenylindole (DAPI) and cover glasses were mounted in ProlongGold (Life Technologies). Images were collected on an inverted epifluorescence microscope (Nikon TE2000E) equipped with a 60x, 1.40 numerical aperture (NA) oil-immersion objective and a 1.5x tube lens on a Photometrics CoolSnapHQ camera driven by NIS-Elements (Nikon) software. Only cells expressing low to medium levels of exogenous motor were selected for imaging.

To prepare cell extracts for fluorescence microtubule-binding assays, COS-7 cells 16 h post-transfection were harvested and centrifuged at low-speed at 4°C. The cell pellet was washed with PBS and resuspended in ice-cold lysis buffer (200 mM NaCl, 4 mM MgCl2, 0.5 mM EDTA, 1 mM EGTA, 0.5% igepal, 7% sucrose, and 20 mM imidazole-HCl, pH 7.5) freshly supplemented with 1 mM ATP, 1 mM phenylmethylsulphonyl fluoride (PMSF) and protease inhibitors cocktail (P8340, Sigma). After centrifugation for 10 min at 20,000 g at 4°C, aliquots of the supernatant were snap frozen in liquid nitrogen and stored at −80 °C until further use. To prepare extracts for microtubule dynamics assays, COS-7 cells 16 h post-transfection were harvested and centrifuged at low-speed at 4°C. The cell pellet was washed with PBS and resuspended in ice-cold BRB80 buffer (80 mM PIPES/KOH, pH 6.8, 1 mM MgCl_2_, 1mM EGTA) freshly supplemented with 1 mM ATP, 1 mM PMSF and protease inhibitors cocktail. The cells were lysed by sonication (Fisher Scientific, Sonic Dismembrator Model 500) using 10% power, 4×10 sec on ice. After centrifugation for 10 min at 20,000 g at 4°C, aliquots of the supernatant were snap frozen in liquid nitrogen and stored at −80 °C until further use.

### Microtubule binding and dynamics assays

For live-cell imaging, COS-7 cells seeded onto glass-bottomed dishes (MatTek Corporation) were co-transfected with plasmids for expression of EB3-mCherry with the indicated mNeonGreen-tagged motors. After ^~^5h, the cells were washed once then incubated in Leibovitz’s L-15 medium (Gibco) and imaged at 37°C in a temperature-controlled and humidified live-imaging chamber (Tokai Hit) mounted on an inverted total internal reflection fluorescence (TIRF) microscope Ti-E/B (Nikon) equipped with a 100× 1.49 N.A. oil immersion TIRF objective (Nikon), three 20 mW diode lasers (488 nm, 561 nm and 640 nm), and EMCCD detector (iXon X3DU897, Andor). The angle of illumination was adjusted for maximum penetration of the evanescent field into the cell. Image acquisition was controlled with Nikon Element software. Images were acquired in both 488- and 561-nm channels every 100 ms for 30 s.

For measuring microtubule dynamics *in vitro*, a flow cell (^~^10 μl volume) was assembled by attaching a clean #1.5 coverslip (Fisher Scientific) to a glass slide (Fisher Scientific) with two strips of double-sided tape. Microtubule seeds containing 10% X-rhodamine-labeled and 10% biotin-labeled tubulin (Cytoskeleton Inc.) were generated by polymerization in the presence of the non-hydrolysable GTP analogue GMPCPP (NU-405S, Jena Bioscience) and then immobilized on coverslips incubated sequentially with the following solutions: (1) 1 mg/ml BSA-biotin (A8549, Sigma), (2) blocking buffer (1 mg/ml BSA in BRB80), (3) 0.5 mg/ml NeutrAvidin (31000, Thermo), (4) blocking buffer, (5) short GMPCPP-stabilized microtubule seeds, (6) blocking buffer. Microtubule growth was then initiated by flowing in 20 μM tubulin containing 7% Hilyte647-labeled tubulin (Cytoskeleton Inc.) together with cell extracts in the reaction buffer [2.5 mM GTP, 2.5 mM ATP, 0.2 mg/ml BSA, 2 mg/ml casein, 2 mM MgCl_2_, 0.2% methy-cellulose (Sigma) and oxygen-scavenging (2 mM DTT, 20 mM glucose, 0.4 mg/ml glucose oxidase, and 0.16 mg/ml catalase) in BRB80]. The flow cells were sealed with molten paraffin wax and imaged by TIRF microscopy. Time-lapse images were acquired in 488nm, 561 nm and 640 nm channels at a rate of every 5 s for 15 min. The temperature was set at 35°C in a temperature-controlled chamber (Tokai Hit). To determine of the growth rate and catastrophe frequency of microtubule plus ends, maximum intensity projections were generated and kymographs (width= 3 pixels) were generated using ImageJ and displayed with time on the y axis and distance on the x axis. Only growth events with a slope over a three-pixel length were analyzed. Catastrophe frequency was obtained by dividing the total number of catastrophes observed by the imaging time (15 min)..

For the microtubule binding assay, Hilyte-647 labeled microtubules were polymerized from purified tubulin (Cytoskeleton Inc.) in BRB80 Buffer supplemented with 2.5 mM GTP and 2 mM MgCl_2_ at 37°C for 30 min. Polymerized microtubules were stored in the dark at room temperature after addition of ten volumes of prewarmed BRB80 containing 20 μM taxol and additional 30 min incubation at 37°C. The amount of motors in COS-7 lysates was determined by Dot Blot using an anti-GFP antibody (1:10,000, ProteinTech, #66002-1). Digital images of the blots were analyzed by ImageJ. Equal amounts of motors in the final motility mixture [6 mg/ml BSA, 0.5 mg/ml casein, 10 μM taxol, 2 mM ATP, 1 mM MgCl_2_ and oxygen-scavenging system in P12 buffer (12 mM Pipes/KOH, pH 6.8, 2 mM MgCl2, and 1 mM EGTA] were added to flow cells containing polymerized microtubules. The coverslips were sealed and imaged at room temperature by TIRF microscopy. All the images were acquired and analyzed using the same conditions. For each motor, the fluorescence intensities along the microtubules were measured using ImageJ and the fluorescence intensity of an adjacent region was subtracted as background. The experiment was repeated two times for each motor with similar results.

## Acknowledgments

We thank members of the Verhey lab for helpful discussions. We thank Takashi Hotta for help with sonicated cell exacts. The work was funded by grants from the National Institutes of Health (R01GM070862, R01GM118751).

## References

1. Hirokawa, N., Noda, Y., Tanaka, Y., and Niwa, S. (2009). Kinesin superfamily motor proteins and intracellular transport. Nat Rev Mol Cell Biol. 10: 682–96.

2. Tomishige, M., Klopfenstein, D.R., and Vale, R.D. (2002). Conversion of Unc104/KIF1A kinesin into a processive motor after dimerization. Science. 297: 2263–7.

3. Cai, D., McEwen, D.P., Martens, J.R., Meyhofer, E., and Verhey, K.J. (2009). Single molecule imaging reveals differences in microtubule track selection between Kinesin motors. PLoS Biol. 7: e1000216.

4. Soppina, V., Norris, S.R., Dizaji, A.S., Kortus, M., Veatch, S., Peckham, M., and Verhey, K.J. (2014). Dimerization of mammalian kinesin-3 motors results in superprocessive motion. Proc Natl Acad Sci U S A. 111: 5562–7.

5. Hoeprich, G.J., Mickolajczyk, K.J., Nelson, S.R., Hancock, W.O., and Berger, C.L. (2017). The axonal transport motor kinesin-2 navigates microtubule obstacles via protofilament switching. Traffic. 18: 304–314.

6. Andreasson, J.O., Shastry, S., Hancock, W.O., and Block, S.M. (2015). The Mechanochemical Cycle of Mammalian Kinesin-2 KIF3A/B under Load. Curr Biol. 25: 1166–75.

7. Guzik-Lendrum, S., Rank, K.C., Bensel, B.M., Taylor, K.C., Rayment, I., and Gilbert, S.P. (2015). Kinesin-2 KIF3AC and KIF3AB Can Drive Long-Range Transport along Microtubules. Biophys J. 109: 1472–82.

8. Cross, R.A. (2016). Review: Mechanochemistry of the kinesin-1 ATPase. Biopolymers. 105: 476–82.

9. Hancock, W.O. (2016). The Kinesin-1 Chemomechanical Cycle: Stepping Toward a Consensus. Biophys J. 110: 1216–25.

10. Feng, Q., Mickolajczyk, K.J., Chen, G.Y., and Hancock, W.O. (2018). Motor Reattachment Kinetics Play a Dominant Role in Multimotor-Driven Cargo Transport. Biophys J. 114: 400–409.

11. Cross, R.A. and McAinsh, A. (2014). Prime movers: the mechanochemistry of mitotic kinesins. Nat Rev Mol Cell Biol. 15: 257–71.

12. Friel, C.T. and Howard, J. (2012). Coupling of kinesin ATP turnover to translocation and microtubule regulation: one engine, many machines. J Muscle Res Cell Motil. 33: 377–83.

13. Friel, C.T. and Welburn, J.P. (2018). Parts list for a microtubule depolymerising kinesin. Biochem Soc Trans. 46: 1665–1672.

14. Su, X., Ohi, R., and Pellman, D. (2012). Move in for the kill: motile microtubule regulators. Trends Cell Biol. 22: 567–75.

15. Mann, B.J. and Wadsworth, P. (2019). Kinesin-5 Regulation and Function in Mitosis. Trends Cell Biol. 29: 66–79.

16. Singh, S.K., Pandey, H., Al-Bassam, J., and Gheber, L. (2018). Bidirectional motility of kinesin-5 motor proteins: structural determinants, cumulative functions and physiological roles. Cell Mol Life Sci. 75: 1757–1771.

17. Endoh-Yamagami, S., Evangelista, M., Wilson, D., Wen, X., Theunissen, J.W., Phamluong, K., Davis, M., Scales, S.J., Solloway, M.J., de Sauvage, F.J., and Peterson, A.S. (2009). The mammalian Cos2 homolog Kif7 plays an essential role in modulating Hh signal transduction during development. Curr Biol. 19: 1320–6.

18. Cheung, H.O., Zhang, X., Ribeiro, A., Mo, R., Makino, S., Puviindran, V., Law, K.K., Briscoe, J., and Hui, C.C. (2009). The kinesin protein Kif7 is a critical regulator of Gli transcription factors in mammalian hedgehog signaling. Sci Signal. 2: ra29.

19. Ali, B.R., Silhavy, J.L., Akawi, N.A., Gleeson, J.G., and Al-Gazali, L. (2012). A mutation in KIF7 is responsible for the autosomal recessive syndrome of macrocephaly, multiple epiphyseal dysplasia and distinctive facial appearance. Orphanet J Rare Dis. 7: 27.

20. Barakeh, D., Faqeih, E., Anazi, S., M, S.A.-D., Softah, A., Albadr, F., Hassan, H., Alazami, A.M., and Alkuraya, F.S. (2015). The many faces of KIF7. Hum Genome Var. 2: 15006.

21. Bourke, J.P., Watson, G., Muntoni, F., Spinty, S., Roper, H., Guglieri, M., Speed, C., McColl, E., Chikermane, A., Jayawant, S., Adwani, S., Willis, T., Wilkinson, J., Bryant, A., Chadwick, T., Wood, R., Bushby, K., and group, D.M.D.H.P.s. (2018). Randomised placebo-controlled trial of combination ACE inhibitor and beta-blocker therapy to prevent cardiomyopathy in children with Duchenne muscular dystrophy? (DMD Heart Protection Study): a protocol study. BMJ Open. 8: e022572.

22. Karaer, K., Yuksel, Z., Ichkou, A., Calisir, C., and Attie-Bitach, T. (2015). A novel KIF7 mutation in two affected siblings with acrocallosal syndrome. Clin Dysmorphol. 24: 61–4.

23. Putoux, A., Thomas, S., Coene, K.L., Davis, E.E., Alanay, Y., Ogur, G., Uz, E., Buzas, D., Gomes, C., Patrier, S., Bennett, C.L., Elkhartoufi, N., Frison, M.H., Rigonnot, L., Joye, N., Pruvost, S., Utine, G.E., Boduroglu, K., Nitschke, P., Fertitta, L., Thauvin-Robinet, C., Munnich, A., Cormier-Daire, V., Hennekam, R., Colin, E., Akarsu, N.A., Bole-Feysot, C., Cagnard, N., Schmitt, A., Goudin, N., Lyonnet, S., Encha-Razavi, F., Siffroi, J.P., Winey, M., Katsanis, N., Gonzales, M., Vekemans, M., Beales, P.L., and Attie-Bitach, T. (2011). KIF7 mutations cause fetal hydrolethalus and acrocallosal syndromes. Nat Genet. 43: 601–6.

24. Tunovic, S., Baranano, K.W., Barkovich, J.A., Strober, J.B., Jamal, L., and Slavotinek, A.M. (2015). Novel KIF7 missense substitutions in two patients presenting with multiple malformations and features of acrocallosal syndrome. Am J Med Genet A. 167A: 2767–76.

25. Walsh, D.M., Shalev, S.A., Simpson, M.A., Morgan, N.V., Gelman-Kohan, Z., Chemke, J., Trembath, R.C., and Maher, E.R. (2013). Acrocallosal syndrome: identification of a novel KIF7 mutation and evidence for oligogenic inheritance. Eur J Med Genet. 56: 39–42.

26. Ibisler, A., Hehr, U., Barth, A., Koch, M., Epplen, J.T., and Hoffjan, S. (2015). Novel KIF7 Mutation in a Tunisian Boy with Acrocallosal Syndrome: Case Report and Review of the Literature. Mol Syndromol. 6: 173–80.

27. Dafinger, C., Liebau, M.C., Elsayed, S.M., Hellenbroich, Y., Boltshauser, E., Korenke, G.C., Fabretti, F., Janecke, A.R., Ebermann, I., Nurnberg, G., Nurnberg, P., Zentgraf, H., Koerber, F., Addicks, K., Elsobky, E., Benzing, T., Schermer, B., and Bolz, H.J. (2011). Mutations in KIF7 link Joubert syndrome with Sonic Hedgehog signaling and microtubule dynamics. J Clin Invest. 121: 2662–7.

28. Asadollahi, R., Strauss, J.E., Zenker, M., Beuing, O., Edvardson, S., Elpeleg, O., Strom, T.M., Joset, P., Niedrist, D., Otte, C., Oneda, B., Boonsawat, P., Azzarello-Burri, S., Bartholdi, D., Papik, M., Zweier, M., Haas, C., Ekici, A.B., Baumer, A., Boltshauser, E., Steindl, K., Nothnagel, M., Schinzel, A., Stoeckli, E.T., and Rauch, A. (2018). Clinical and experimental evidence suggest a link between KIF7 and C5orf42-related ciliopathies through Sonic Hedgehog signaling. Eur J Hum Genet. 26: 197–209.

29. Coles, G.L., Baglia, L.A., and Ackerman, K.G. (2015). KIF7 Controls the Proliferation of Cells of the Respiratory Airway through Distinct Microtubule Dependent Mechanisms. PLoS Genet. 11: e1005525.

30. Wong, K.Y., Liu, J., and Chan, K.W. (2017). KIF7 attenuates prostate tumor growth through LKB1-mediated AKT inhibition. Oncotarget. 8: 54558–54571.

31. Lau, C.I., Barbarulo, A., Solanki, A., Saldana, J.I., and Crompton, T. (2017). The kinesin motor protein Kif7 is required for T-cell development and normal MHC expression on thymic epithelial cells (TEC) in the thymus. Oncotarget. 8: 24163–24176.

32. Liem, K.F., Jr., He, M., Ocbina, P.J., and Anderson, K.V. (2009). Mouse Kif7/Costal2 is a cilia-associated protein that regulates Sonic hedgehog signaling. Proc Natl Acad Sci U S A. 106: 13377–82.

33. He, M., Subramanian, R., Bangs, F., Omelchenko, T., Liem, K.F., Jr., Kapoor, T.M., and Anderson, K.V. (2014). The kinesin-4 protein Kif7 regulates mammalian Hedgehog signalling by organizing the cilium tip compartment. Nat Cell Biol. 16: 663–72.

34. Emechebe, U., Kumar, P.P., Rozenberg, J.M., Moore, B., Firment, A., Mirshahi, T., and Moon, A.M. (2016). T-box3 is a ciliary protein and regulates stability of the Gli3 transcription factor to control digit number. Elife. 5.

35. Putoux, A., Alqahtani, A., Pinson, L., Paulussen, A.D., Michel, J., Besson, A., Mazoyer, S., Borg, I., Nampoothiri, S., Vasiljevic, A., Uwineza, A., Boggio, D., Champion, F., de Die-Smulders, C.E., Gardeitchik, T., van Putten, W.K., Perez, M.J., Musizzano, Y., Razavi, F., Drunat, S., Verloes, A., Hennekam, R., Guibaud, L., Alix, E., Sanlaville, D., Lesca, G., and Edery, P. (2016). Refining the phenotypical and mutational spectrum of Taybi-Linder syndrome. Clin Genet. 90: 550–555.

36. Jiang, S., Mani, N., Wilson-Kubalek, E.M., Ku, P.I., Milligan, R.A., and Subramanian, R. (2019). Interplay between the Kinesin and Tubulin Mechanochemical Cycles Underlies Microtubule Tip Tracking by the Non-motile Ciliary Kinesin Kif7. Dev Cell. 49: 711–730 e8.

37. Yue, Y., Blasius, T.L., Zhang, S., Jariwala, S., Walker, B., Grant, B.J., Cochran, J.C., and Verhey, K.J. (2018). Altered chemomechanical coupling causes impaired motility of the kinesin-4 motors KIF27 and KIF7. J Cell Biol. 217: 1319–1334.

38. Verhey, K.J. and Hammond, J.W. (2009). Traffic control: regulation of kinesin motors. Nat Rev Mol Cell Biol. 10: 765–77.

39. Coles, G.L. and Ackerman, K.G. (2013). Kif7 is required for the patterning and differentiation of the diaphragm in a model of syndromic congenital diaphragmatic hernia. Proc Natl Acad Sci U S A. 110: E1898–905.

40. Maurya, A.K., Ben, J., Zhao, Z., Lee, R.T., Niah, W., Ng, A.S., Iyu, A., Yu, W., Elworthy, S., van Eeden, F.J., and Ingham, P.W. (2013). Positive and negative regulation of Gli activity by Kif7 in the zebrafish embryo. PLoS Genet. 9: e1003955.

41. Hammond, J.W., Blasius, T.L., Soppina, V., Cai, D., and Verhey, K.J. (2010). Autoinhibition of the kinesin-2 motor KIF17 via dual intramolecular mechanisms. J Cell Biol. 189: 1013–25.

42. Hammond, J.W., Cai, D., Blasius, T.L., Li, Z., Jiang, Y., Jih, G.T., Meyhofer, E., and Verhey, K.J. (2009). Mammalian Kinesin-3 motors are dimeric in vivo and move by processive motility upon release of autoinhibition. PLoS Biol. 7: e72.

43. Nakata, T. and Hirokawa, N. (2003). Microtubules provide directional cues for polarized axonal transport through interaction with kinesin motor head. J Cell Biol. 162: 1045–55.

44. Nakata, T. and Hirokawa, N. (1995). Point mutation of adenosine triphosphate-binding motif generated rigor kinesin that selectively blocks anterograde lysosome membrane transport. J Cell Biol. 131: 1039–53.

45. Klejnot, M. and Kozielski, F. (2012). Structural insights into human Kif7, a kinesin involved in Hedgehog signalling. Acta Crystallogr D Biol Crystallogr. 68: 154–9.

46. Shastry, S. and Hancock, W.O. (2010). Neck linker length determines the degree of processivity in kinesin-1 and kinesin-2 motors. Curr Biol. 20: 939–43.

47. Shastry, S. and Hancock, W.O. (2011). Interhead tension determines processivity across diverse N-terminal kinesins. Proc Natl Acad Sci U S A. 108: 16253–8.

48. Cheng, L., Desai, J., Miranda, C.J., Duncan, J.S., Qiu, W., Nugent, A.A., Kolpak, A.L., Wu, C.C., Drokhlyansky, E., Delisle, M.M., Chan, W.M., Wei, Y., Propst, F., Reck-Peterson, S.L., Fritzsch, B., and Engle, E.C. (2014). Human CFEOM1 mutations attenuate KIF21A autoinhibition and cause oculomotor axon stalling. Neuron. 82: 334–49.

49. Bianchi, S., van Riel, W.E., Kraatz, S.H., Olieric, N., Frey, D., Katrukha, E.A., Jaussi, R., Missimer, J., Grigoriev, I., Olieric, V., Benoit, R.M., Steinmetz, M.O., Akhmanova, A., and Kammerer, R.A. (2016). Structural basis for misregulation of kinesin KIF21A autoinhibition by CFEOM1 disease mutations. Sci Rep. 6: 30668.

50. van der Vaart, B., van Riel, W.E., Doodhi, H., Kevenaar, J.T., Katrukha, E.A., Gumy, L., Bouchet, B.P., Grigoriev, I., Spangler, S.A., Yu, K.L., Wulf, P.S., Wu, J., Lansbergen, G., van Battum, E.Y., Pasterkamp, R.J., Mimori-Kiyosue, Y., Demmers, J., Olieric, N., Maly, I.V., Hoogenraad, C.C., and Akhmanova, A. (2013). CFEOM1-associated kinesin KIF21A is a cortical microtubule growth inhibitor. Dev Cell. 27: 145–60.

51. van Riel, W.E., Rai, A., Bianchi, S., Katrukha, E.A., Liu, Q., Heck, A.J., Hoogenraad, C.C., Steinmetz, M.O., Kapitein, L.C., and Akhmanova, A. (2017). Kinesin-4 KIF21B is a potent microtubule pausing factor. Elife. 6: e24746.

52. Matthies, H.J., Baskin, R.J., and Hawley, R.S. (2001). Orphan kinesin NOD lacks motile properties but does possess a microtubule-stimulated ATPase activity. Mol Biol Cell. 12: 4000–12.

53. Bieling, P., Telley, I.A., and Surrey, T. (2010). A minimal midzone protein module controls formation and length of antiparallel microtubule overlaps. Cell. 142: 420–32.

54. Ghiretti, A.E., Thies, E., Tokito, M.K., Lin, T., Ostap, E.M., Kneussel, M., and Holzbaur, E.L.F. (2016). Activity-Dependent Regulation of Distinct Transport and Cytoskeletal Remodeling Functions of the Dendritic Kinesin KIF21B. Neuron. 92: 857–872.

55. Hu, C.K., Coughlin, M., Field, C.M., and Mitchison, T.J. (2011). KIF4 regulates midzone length during cytokinesis. Curr Biol. 21: 815–24.

56. Stumpff, J., Wagenbach, M., Franck, A., Asbury, C.L., and Wordeman, L. (2012). Kif18A and chromokinesins confine centromere movements via microtubule growth suppression and spatial control of kinetochore tension. Dev Cell. 22: 1017–29.

57. Wandke, C., Barisic, M., Sigl, R., Rauch, V., Wolf, F., Amaro, A.C., Tan, C.H., Pereira, A.J., Kutay, U., Maiato, H., Meraldi, P., and Geley, S. (2012). Human chromokinesins promote chromosome congression and spindle microtubule dynamics during mitosis. J Cell Biol. 198: 847–63.

58. Nunes Bastos, R., Gandhi, S.R., Baron, R.D., Gruneberg, U., Nigg, E.A., and Barr, F.A. (2013). Aurora B suppresses microtubule dynamics and limits central spindle size by locally activating KIF4A. J Cell Biol. 202: 605–21.

59. Muhia, M., Thies, E., Labonte, D., Ghiretti, A.E., Gromova, K.V., Xompero, F., Lappe-Siefke, C., Hermans-Borgmeyer, I., Kuhl, D., Schweizer, M., Ohana, O., Schwarz, J.R., Holzbaur, E.L.F., and Kneussel, M. (2016). The Kinesin KIF21B Regulates Microtubule Dynamics and Is Essential for Neuronal Morphology, Synapse Function, and Learning and Memory. Cell Rep. 15: 968–977.

60. Budaitis, B.G., Jariwala, S., Reinemann, D.N., Schimert, K.I., Scarabelli, G., Grant, B.J., Sept, D., Lang, M.J., and Verhey, K.J. (2019). Neck linker docking is critical for Kinesin-1 force generation in cells but at a cost to motor speed and processivity. Elife. 8.

61. Yildiz, A., Tomishige, M., Gennerich, A., and Vale, R.D. (2008). Intramolecular strain coordinates kinesin stepping behavior along microtubules. Cell. 134: 1030–41.

62. Gicking, A.M., Wang, P., Liu, C., Mickolajczyk, K.J., Guo, L., Hancock, W.O., and Qiu, W. (2019). The Orphan Kinesin PAKRP2 Achieves Processive Motility via a Noncanonical Stepping Mechanism. Biophys J. 116: 1270–1281.

63. Guan, R., Zhang, L., Su, Q.P., Mickolajczyk, K.J., Chen, G.Y., Hancock, W.O., Sun, Y., Zhao, Y., and Chen, Z. (2017). Crystal structure of Zen4 in the apo state reveals a missing conformation of kinesin. Nat Commun. 8: 14951.

64. Walker, B.C., Tempel, W., Zhu, H., Park, H., and Cochran, J.C. (2019). Chromokinesins NOD and KID Use Distinct ATPase Mechanisms and Microtubule Interactions To Perform a Similar Function. Biochemistry. 58: 2326–2338.

65. Hutterer, A., Glotzer, M., and Mishima, M. (2009). Clustering of centralspindlin is essential for its accumulation to the central spindle and the midbody. Curr Biol. 19: 2043–9.

66. Yajima, J., Edamatsu, M., Watai-Nishii, J., Tokai-Nishizumi, N., Yamamoto, T., and Toyoshima, Y.Y. (2003). The human chromokinesin Kid is a plus end-directed microtubule-based motor. EMBO J. 22: 1067–74.

67. Li, C., Xue, C., Yang, Q., Low, B.C., and Liou, Y.C. (2016). NuSAP governs chromosome oscillation by facilitating the Kid-generated polar ejection force. Nat Commun. 7: 10597.

68. Davies, T., Kodera, N., Kaminski Schierle, G.S., Rees, E., Erdelyi, M., Kaminski, C.F., Ando, T., and Mishima, M. (2015). CYK4 promotes antiparallel microtubule bundling by optimizing MKLP1 neck conformation. PLoS Biol. 13: e1002121.

69. White, E.A., Raghuraman, H., Perozo, E., and Glotzer, M. (2013). Binding of the CYK-4 subunit of the centralspindlin complex induces a large scale conformational change in the kinesin subunit. J Biol Chem. 288: 19785–95.

70. Tao, L., Fasulo, B., Warecki, B., and Sullivan, W. (2016). Tum/RacGAP functions as a switch activating the Pav/kinesin-6 motor. Nat Commun. 7: 11182.

71. Chong, Y.C., Mann, R.K., Zhao, C., Kato, M., and Beachy, P.A. (2015). Bifurcating action of Smoothened in Hedgehog signaling is mediated by Dlg5. Genes Dev. 29: 262–76.

